# 1.4 GHz amplitude-modulated radiofrequency electromagnetic waves effects on neural cultures

**DOI:** 10.1101/2025.10.10.680616

**Authors:** Gabriel Gaugain, Anne Canovi, Rosa Orlacchio, Florence Poulletier de Gannes, Denys Nikolayev, Julien Modolo

## Abstract

Non-invasive brain stimulation (NIBS) technologies have the potential to positively impact the treatment of neurological disorders. However, NIBS techniques suffer from a lack of penetration depth and focality, thereby restraining their full potential. Here, we investigate the feasibility of a novel technique based on sinusoidal radiofrequency electromagnetic fields (RF-EMF) that are amplitude-modulated in the extremely low frequency (ELF) range. To assess the neuromodulatory effects of ELF-modulated RF-EMF, we exposed primary neuronal cultures placed on Multi-Electrode Arrays (MEAs) and quantified their spiking activity based on phase entrainment. Our results failed to reveal consistent phase entrainment across cultures, while replicating a decrease of neuronal activity with purely sinusoidal 1.4 GHz stimulation. Finally, we suggest protocol improvements that might result in more reproducible effects, which is indispensable for translational applications.

## INTRODUCTION

Stimulation of the brain with electric or magnetic fields has generated considerable interest within the last 40 years for the treatment of neurological disorders. One of the most well-known examples of successful brain stimulation therapies is deep brain stimulation (DBS) in Parkinson’s disease, used for the symptomatic treatment of tremor [1]. Despite its wide use and high efficacy, this technique suffers from drawbacks, and notably invasiveness due to the electrodes that need to be implanted surgically in deep brain regions (e.g., subthalamic nucleus [2]). Therefore, this has motivated the development of brain stimulation techniques that are non-invasive (not requiring surgery), such as transcranial direct / alternating current stimulation (tDCS / tACS), which uses electrodes placed on the scalp to deliver currents up to 2 mA with the objective to achieve modulation of the underlying cortical tissue. However, despite their growing recognized efficacy, for example, in improving cognitive functions such as working memory [3], [4] or reducing the symptoms of major depressive disorder [5], tDCS / tACS are limited in terms of focality and level of electric field that can be achieved at the cortical tissue level, due to significant current shunting (e.g., by the cerebrospinal fluid).

Here, to overcome those drawbacks, we investigated *in vitro* the feasibility of a novel approach for non-invasive, focal brain stimulation using a radiofrequency (RF) sinusoidal waveform modulated in the extremely low-frequency (ELF) range. The rationale for this choice is the following: RF fields (for example, in the 1-3 GHz range), due to their short wavelength, offer the potential for spatial focality. However, the membrane of neurons is supposedly not sensitive to depolarizations within the frequency range of RF, which far exceed the time scale of the ionic channels that underlie membrane polarization, and also their maximal firing rates (typically on the order of 100 Hz). Therefore, we hypothesize that an RF field amplitude-modulated in the ELF range can effectively influence Central Nervous System (CNS) activity, enabling modulatory effects at frequencies capable of producing physiologically meaningful effects. This hypothesis was supported by recent developments using kHz range AM stimuli known as temporal interference (TI, [6]) as well as older findings using very high frequency carriers [7], [8]. These ideas rely on the ability of neurons to rectify electrical stimulus [9], [10], [11]. Using even higher frequencies could, in principle, enable focal stimulation of the CNS at frequencies that are relevant for electrophysiological activities.

To achieve this objective, we used primary neuronal cultures placed on multi-electrode arrays (MEA) to record electrophysiological activity of neurons during exposure to ELF-modulated RF fields. If our hypothesis is correct, and that ELF modulation frequency impacts neuronal activity, this should result in a measurable phase entrainment between the ELF modulation waveform and the neuronal spikes recorded in the electrophysiological signals. We tested a total of N=7 neuronal cultures, applying a 1.4 GHz RF field-chosen for its optimized tissue penetration [12]-with a 2 Hz modulation frequency, which is within the range of spontaneous electrophysiological activity recorded in neuronal cultures [13]. The originality of our approach is the use of sinusoidal low-frequency modulation, rather than the pulsed modulation commonly used in commercial RF systems, as electrophysiological activity itself is typically sinusoidal.

## METHODS

### Primary culture of neurons

We used primary neuronal cultures obtained by the resection of cortical tissue in embryonic (E18-E19) Sprague-Dawley rats as described in Canovi et al. (2023). Briefly, once dissected, embryonic cortices were enzymatically (papain/DNase) and mechanically dissociated, followed by centrifugation through an albumin-inhibitor gradient. The resulting cortical cells (neurons and glia) were resuspended in Neurobasal medium supplemented with 2% B-27, 1% GlutaMAX, and 1% penicillin-streptomycin. Approximately 10^5^ cells were placed onto polylysine/laminin-coated multi-electrode arrays (MEA) containing 60 titanium nitride electrodes (200 μm spaced with 30 μm-diameter tips, 60MEA200/30iR-Ti-Upside Down, Multi-Channel Systems,MCS, GmbH, Reutlingen, Germany). Neurons survival and connectivity prior to data acquisition was verified through microscope examination, to ensure quality and viability of the cultures. Neuron cultures were maintained in an incubator at 37°C, 5% CO2, with the culture medium exchanged every 48 hours until recordings. Neuronal networks were considered mature and were used between 18 and 20 days *in vitro* (DIV) (N= 7).

### Data acquisition

As previously described [14], [15], [16] and schematically illustrated in Figure 1A, the exposure setup for electrophysiology recordings comprised the MEA containing neuronal cultures and an open Transverse ElectroMagnetic (TEM) cell enabling uniform signal propagation through the cell culture, all placed in an incubator maintaining physiological conditions of the cells (37°C, 5% CO2). The TEM cell was directly connected to an RF generation unit, placed outside the incubator (Figure 1A). The RF generation unit comprised (1) an RF generator (R&S®SML02, Rohde & Schwarz, Munich, Germany) producing continuous wave (CW) or 2 Hz-modulated signals at 1.4 GHz with an RF power density of 36 dBm, (2) a 25 dB gain amplifier (ZHL-4W-422+, Mini-circuit, NY, United States), (3) a directional coupler (ZGBDC30-372HP+, Mini-circuit, NY, United States) and (4) a power meter (N1912A, Keysight, United States) equipped with two power sensors for real-time monitoring of incident and reflected powers. Electrophysiological signals were acquired at 10 kHz/channel using the MC Rack software (MCS GmbH), through the connection between the MEA and the pre-amplifier (MEA1060-Inv, MCS GmbH, gain of 1,200), as depicted in Figure 1A. Temperature measurements of the culture medium were performed using a fiber-optic thermometer (Luxtron One, Lumasense Technologies, CA, United States, accuracy ± 5%) to ensure that RF exposure did not raise the culture temperature by more than 1°C, avoiding thermal effects.

**Figure 1.**
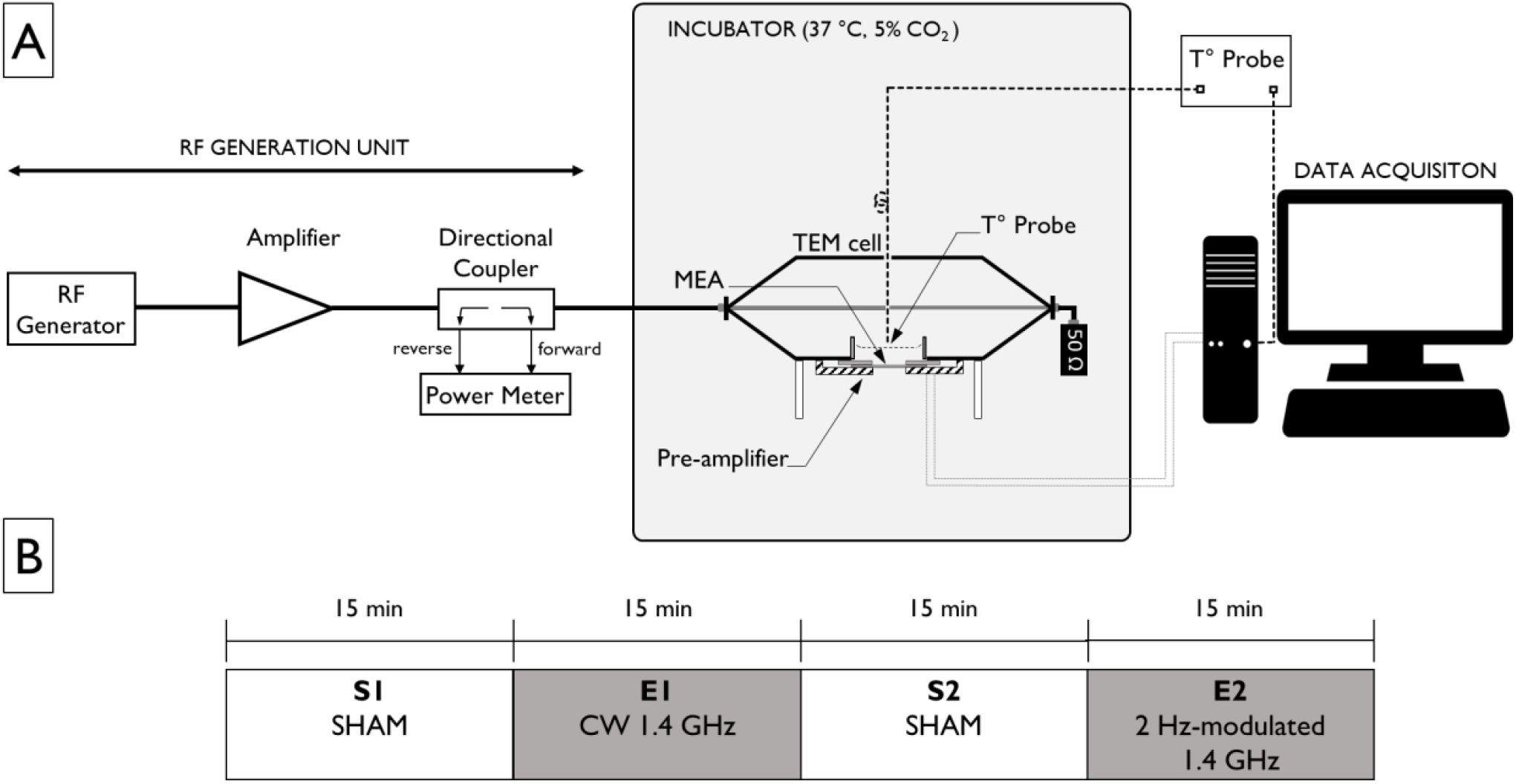
**A) Experimental setup**. The RF generator was coupled to an amplifier to deliver exposure within the incubator in which the TEM cell containing the neuronal culture was placed. A power meter connected to the amplifier *via* a directional coupler was used to monitor incident power in real time during exposure. **B) Exposure protocol**. Temporal course of the exposure, which involved two “Sham” epochs without any stimulation (S1 and S2), and two Exposure epochs (E1 and E2). E1 and E2 both involved exposure to 1.4 GHz RF, but amplitude modulation at 2 Hz was only present during the E2 epoch. Each epoch lasted 15 minutes.

### Exposure Protocol

The exposure protocol consisted of one one-hour period, divided into 4 consecutive blocks of 15 minutes each, as illustrated in Figure 1B. During the first block (S1), no exposure was delivered to the culture (Sham). In the second block (S2), the culture was exposed to a 1.4 GHz continuous, sinusoidal signal at 36 dBm. The third block (S3) again involved no exposure (Sham). Finally, in the fourth block (S4), the culture was exposed to 1.4 GHz (continuous, sinusoidal) signal at 36 dBm, with an additional 2 Hz sinusoidal amplitude modulation. As aforementioned, the choice of 2 Hz as a modulation frequency was made based on the typical low-frequency activity (e.g., bursts) taking place in neuron cultures (the objective being to modulate endogenous activity, which requires a stimulation frequency close to the endogenous frequency). The absence of low-frequency modulation during E1 was designed specifically to identify the specificity of neural entrainment, if any, as a consequence of the 2 Hz modulation only, which is present during E2. Electrophysiological signals were continuously acquired during the exposure protocol.

### Data processing

Electrophysiological signals were read and processed using the [spikeinterface](https://spikeinterface.readthedocs.io/en/stable/) Python package [17]. First, *spikeinterface* was used to perform band-pass filtering between 300 Hz and 4500 Hz as a pre-processing. Second, for each of the 60 channels (electrodes), spikes were detected using a modified version of the differential threshold precision timing spike detection (PTSD) method [18]. Then, in order to test entrainment to the lower-frequency component of the AM-RF exposure, we computed the phase of each spike corresponding to the phase of a 2 Hz sinusoid waveform, independently from the real phase of the signal (preferred phase one phase away), related to the amplitude of the sinusoidal modulation during E2. In the case of Sham epochs, the calculated PLV and “entrainment” exposure is therefore entrainment *per se* (since no stimulation signal is present), but rather represents a locked endogenous rhythm. Polar plots were computed, and the significance of entrainment at a given phase was tested using the Rayleigh test. For all cultures, the mean firing rate (MFR) was computed as in [13]:

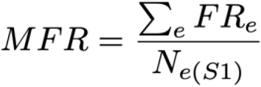

Where FR_e_ is the firing rate of the e^th^ active electrode, and N_e(S1)_ is the number of active electrodes in the S1 phase. To account for variability in basal electrical activity between the different cultures, the MFR was normalized by setting the S1 phase to 100%, such that the MFR reflects changes in firing rate relative to baseline. We used a Friedman test followed by Conover’s multiple comparison test to compare changes in MFR across the different epochs of the exposure protocols.

## RESULTS

After 15 minutes of 20 dBm exposure, the temperature did not increase more than 0.2 °C (not shown). Using this result, it follows that a 15 minute 36 dBm (3.9 W) exposure did not reach the threshold of 1 °C increase (ICNIRP guidelines). Therefore, any reported effects are unlikely to be thermal in origin. Over the 7 cultures recorded with the aforementioned protocol, all had at least 19 electrodes with spiking activity over 0.1 Hz during S1 (see Table 1). Only electrodes reaching this threshold were kept for further analysis to ensure reliable estimations. Specifically, after spike detection, phase entrainment to a 2 Hz sinusoidal waveform was computed to test if AM-RF signal could modulate neural activity in a phase-dependent manner similarly to tACS. Figure 2B presents an example of an electrophysiological signal recorded during exposure. An increase of PLV in the number of entrained electrodes between S1 and E2, as depicted in Figure 2.C, was found (1 to 24 channels (7.8% on average) with an increase in Rayleigh test p-values between E2 and other exposure protocols), showing little effects of the low frequency component on the level of entrainment. Most electrodes had a significant baseline 2 Hz phase synchronization (5 to 42 channels, with a 25% mean), indicative of endogenous 2 Hz activity, as depicted Figure 2.D.

**Figure 2.**
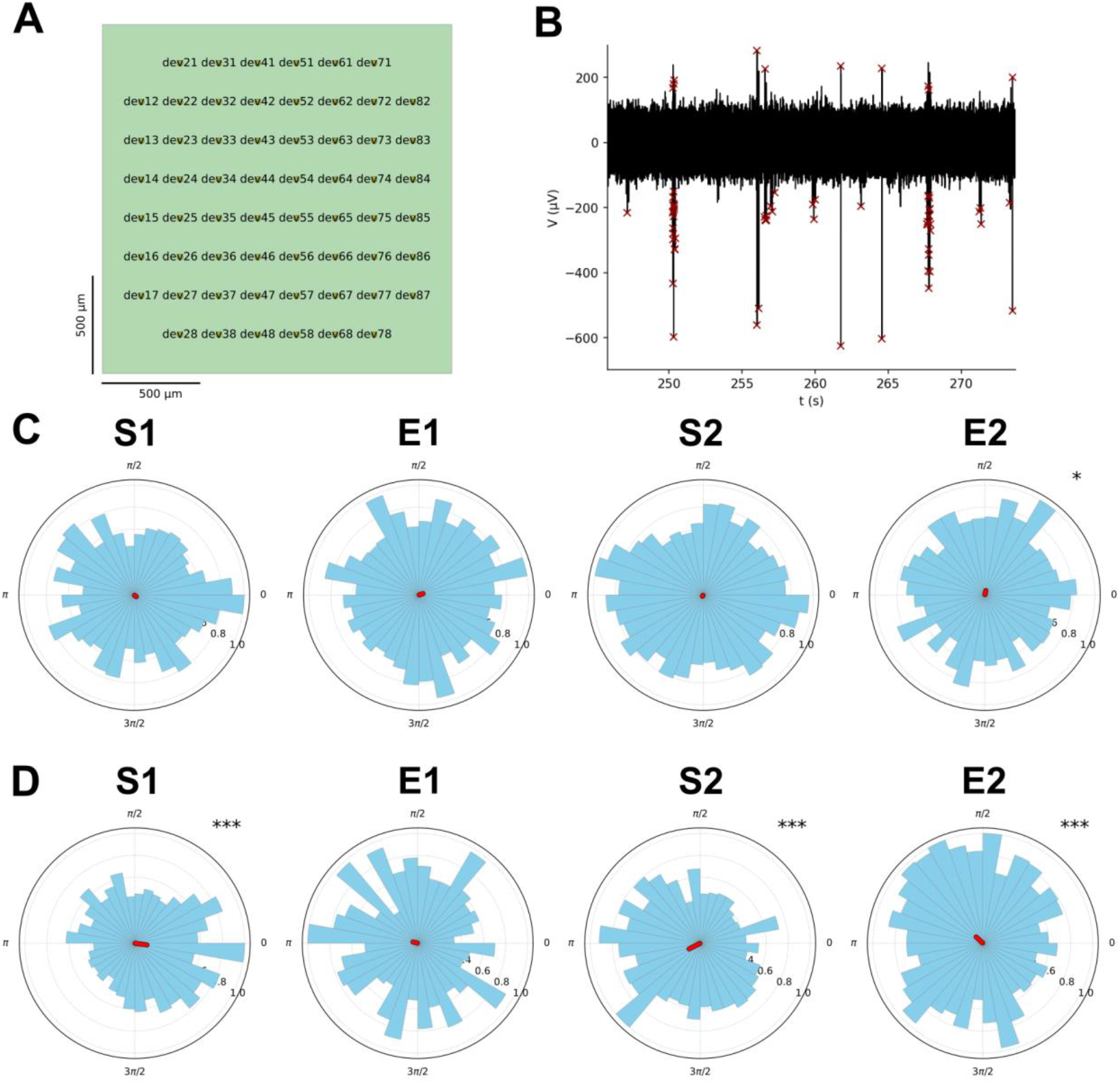
Phase entrainment quantification. **A**: Multi-electrode array (MEA) unit used to record neural activity, with the 60 electrode contacts depicted with their respective id. **B:** Example of a specific voltage trace recorded, each red cross corresponding to a detected spike. **C:** Corresponding polar plots, which indicate the proportion of spikes detected as a specific phase. These polar plots represent action potential phase relative to a 2 Hz sine wave for Sham, RF and AM-RF conditions (in this example, significant phase entrainment only in E2). **D:** Polar plots of the most represented case where phase entrainment was present in multiple protocols, denoting a baseline phase synchrony at 2 Hz endogenous activity. The significance of the Rayleigh test are denoted by: *: p<0.05, **:p<0.01, ***:p<0.001.

The MFR for exposure protocols were compared from baseline, as depicted in Figure 3, and highlighted an activity decrease during 36 dBM sinusoidal RF exposure, while an increase was observed during and after AM-RF exposure.

**Figure 3.**
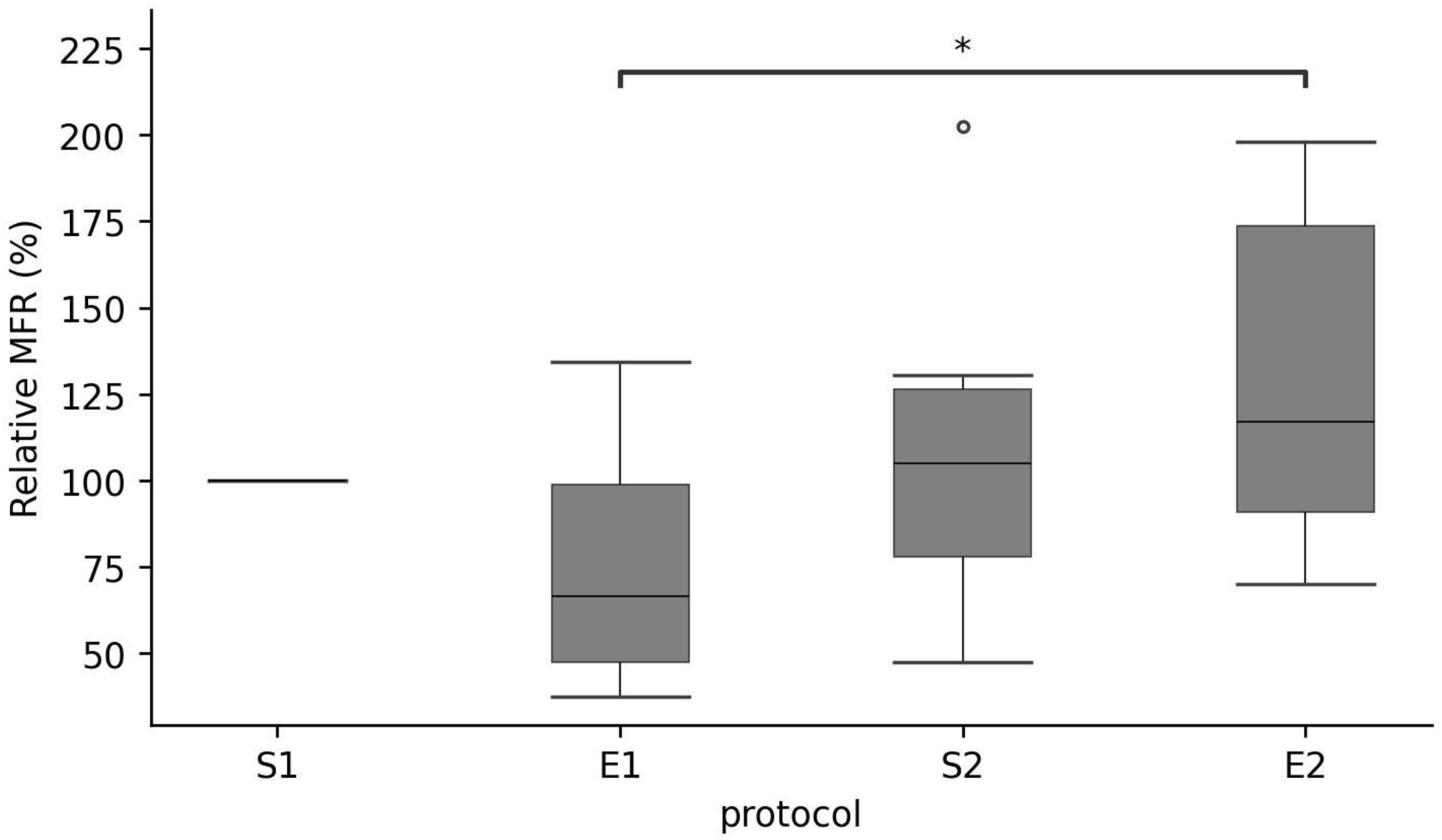
Relative mean firing rates of active electrodes for all cultures and across protocols. The baseline mean firing rate was taken as the S1 being the SHAM condition (100%). A non-significant decrease is visible in E1, while an increase is visible in E2 with significant increase between E1 and E2 (p<0.05, Conover’s multiple comparison test).

## DISCUSSION AND CONCLUDING REMARKS

Here, we aimed at using amplitude-modulated (AM) RF exposure of neuronal cultures to investigate if a low frequency modulation could induce specific neuromodulatory effects. Electrophysiological recordings were performed and associated phase entrainment and spiking activity were assessed before RF exposure, during continuous 1.4 GHz exposure, and with a 2 Hz AM-RF exposure. Our results do not support an effect of low frequency AM on phase entrainment, since few electrodes showed a switch from no entrainment to significant 2 Hz entrainment. However, in terms of firing rate, the results highlighted a decrease of activity during continuous wave 1.4 GHz exposure, in support of previous findings [14], [13].

The firing rate decrease during CW exposure is consistent with the results from several past studies on RF exposure on neuronal cultures [13], [19]. However, the marginal effect of AM-RF exposure contrasts with anterior findings [7], [8] but not with more recent results using metal recording devices [20]. These recent results demonstrated neural entrainment to 5, 10 and 20 Hz AM-RF exposure at 1.8 GHz in mice, but only during a 6.5W exposure, which was above the power used in the present study [20]. However, these results were obtained using metal recordings devices and were not reproduced using a metal-free Ca^2+^ imaging device. In terms of limitations, we acknowledge the possibility of artifacts being due to the interaction of the exposure signal with the MEA electronics, including the electrodes themselves. That being said, we are confident that this possibility of an interaction between the AM-modulated exposure and the electronics as the source of the observed effects is unlikely, since the frequency range of the potential artifact (2 Hz) is very different from the frequency range used for spike quantification (> 500 Hz). Therefore, the quantification of spikes should not be biased by RF modulation artefacts. Another limit is that a neuromodulatory effect could only be detected in one out of seven cultures, therefore our results do not support our initial hypothesis that RF amplitude-modulated in the ELF range (here, 2 Hz) modulates the activity in neuronal cultures. Possible causes include variability in network configuration between neuronal cultures, and insufficient electric field induced *in situ* from the unfavorable orientation of the electric field with respect to neuronal elements. The induced EF is indeed orthogonal to the MEA surface, while neuronal elements such as axons are roughly parallel to it, which according to the widely used “lambda.E” model [21] results in a very low polarization. Alternative configurations of our experimental setup might provide a (challenging) solution to investigate this possibility. A detailed micro- and macro-dosimetric study might also be useful to guide such setup improvements. Importantly, the RF waveforms investigated here are drastically different from those resulting from mobile phone exposure, therefore it is not possible to translate the potential neuromodulatory effect evidenced here with neuromodulatory effects of 4G/5G exposure. Mobile phone RF are indeed more complex, pulsed, with a different frequency composition than those used in our study. Furthermore, this work could benefit from an exhaustive study with counter-balanced order for CW/modulated exposure to avoid possible interactions between E1 (CW 1.4 GHz) and E2 (2 Hz-modulated 1.4 GHz).

In terms of future prospects, we aim at improving our experimental setup to optimize electric field orientation with respect to the MEA surface, which might result in increased entrainment at the AM frequency. While our approach needs further refinement and development, the potential is high for the development of novel focal, non-invasive neurotechnologies that are critically needed with the continuous rise of neurological disorders.

## ACKNOWLEDGMENTS

We acknowledge support of the SP-STIM project by the Labex COMINLABS (No. ANR-10-LABX-07-01).

## REFERENCES

[1] L. Benabid, P. Pollak, A. Louveau, S. Henry, et J. de Rougemont, « Combined (Thalamotomy and Stimulation) Stereotactic Surgery of the VIM Thalamic Nucleus for Bilateral Parkinson Disease », Appl. Neurophysiol., vol. 50, no 1-6, p. 344–346, janv. 1988, doi: 10.1159/000100803.

[2] D. Guehl et al., « Statistical determination of the optimal subthalamic nucleus stimulation site in patients with Parkinson disease », J. Neurosurg., vol. 106, no 1, p. 101–110, janv. 2007, doi: 10.3171/jns.2007.106.1.101.

[3] I. Alekseichuk, Z. Turi, G. Amador de Lara, A. Antal, et W. Paulus, « Spatial working memory in humans fepends on theta and high gamma synchronization in the prefrontal cortex », Curr. Biol., vol. 26, no 12, p. 1513–1521, juin 2016, doi: 10.1016/j.cub.2016.04.035.

[4] R. M. G. Reinhart et J. A. Nguyen, « Working memory revived in older adults by synchronizing rhythmic brain circuits », Nat. Neurosci., vol. 22, no 5, p. 820–827, 2019, doi: 10.1038/s41593-019-0371-x.

[5] J. Zhou et al., « Effect of add-on transcranial alternating current stimulation (tACS) in major depressive disorder: A randomized controlled trial », Brain Stimulat., vol. 17, no 4, p. 760–768, juill. 2024, doi: 10.1016/j.brs.2024.06.004.

[6] N. Grossman et al., « Noninvasive deep brain stimulation via temporally interfering electric fields », Cell, vol. 169, no 6, p. 1029–1041.e16, juin 2017, doi: 10.1016/j.cell.2017.05.024.

[7] S. M. Bawin, R. J. Gavalas-Medici, et W. R. Adey, « Effects of modulated very high frequency fields on specific brain rhythms in cats », Brain Res., vol. 58, no 2, p. 365–384, août 1973, doi: 10.1016/0006-8993(73)90008-5.

[8] R. C. Beason et P. Semm, « Responses of neurons to an amplitude modulated microwave stimulus », Neurosci. Lett., vol. 333, no 3, p. 175–178, nov. 2002, doi: 10.1016/S0304-3940(02)00903-5.

[9] D. E. Goldman, « Potential, impedance, and rectification in membranes », J. Gen. Physiol., vol. 27, no 1, p. 37–60, sept. 1943, doi: 10.1085/jgp.27.1.37.

[10] K. S. Cole, « RECTIFICATION AND INDUCTANCE IN THE SQUID GIANT AXON », J. Gen. Physiol., vol. 25, no 1, p. 29–51, sept. 1941, doi: 10.1085/jgp.25.1.29.

[11] E. Mirzakhalili, B. Barra, M. Capogrosso, et S. F. Lempka, « Biophysics of temporal interference stimulation », Cell Syst., vol. 11, no 6, p. 557–572.e5, déc. 2020, doi: 10.1016/j.cels.2020.10.004.

[12] D. Nikolayev, W. Joseph, M. Zhadobov, R. Sauleau, et L. Martens, « Optimal Radiation of Body-Implanted Capsules », Phys. Rev. Lett., vol. 122, no 10, p. 108101, mars 2019, doi: 10.1103/PhysRevLett.122.108101.

[13] A. Canovi et al., « In vitro exposure of neuronal networks to the 5G-3.5 GHz signal », Front. Public Health, vol. 11, août 2023, doi: 10.3389/fpubh.2023.1231360.

[14] D. Moretti et al., « In-vitro exposure of neuronal networks to the GSM-1800 signal », Bioelectromagnetics, vol. 34, no 8, p. 571–578, 2013, doi: 10.1002/bem.21805.

[15] El Khoueiry et al., « Decreased spontaneous electrical activity in neuronal networks exposed to radiofrequency 1,800 MHz signals », J. Neurophysiol., vol. 120, no 6, p. 2719–2729, déc. 2018, doi: 10.1152/jn.00589.2017.

[16] A. Nefzi et al., « Dosimetry of Microelectrodes Array Chips for Electrophysiological Studies Under Simultaneous Radio Frequency Exposures », IEEE Trans. Microw. Theory Tech., vol. 70, no 3, p. 1871–1881, mars 2022, doi: 10.1109/TMTT.2021.3136296.

[17] P. Buccino et al., « SpikeInterface, a unified framework for spike sorting », eLife, vol. 9, p. e61834, nov. 2020, doi: 10.7554/eLife.61834.

[18] Maccione, M. Gandolfo, P. Massobrio, A. Novellino, S. Martinoia, et M. Chiappalone, « A novel algorithm for precise identification of spikes in extracellularly recorded neuronal signals », J. Neurosci. Methods, vol. 177, no 1, p. 241–249, févr. 2009, doi: 10.1016/j.jneumeth.2008.09.026.

[19] El Khoueiry et al., « Decreased spontaneous electrical activity in neuronal networks exposed to radiofrequency 1,800 MHz signals », J. Neurophysiol., vol. 120, no 6, p. 2719–2729, déc. 2018, doi: 10.1152/jn.00589.2017.

[20] I. Yaghmazadeh et al., « Neuronal activity under transcranial radio-frequency stimulation in metal-free rodent brains in-vivo », Commun. Eng., vol. 1, no 1, Art. no 1, juill. 2022, doi: 10.1038/s44172-022-00014-7.

[21] G. Ruffini et al., « Transcranial current brain stimulation (tCS): Models and technologies », IEEE Trans. Neural Syst. Rehabil. Eng., vol. 21, no 3, p. 333–345, 2013-5 2013, doi: 10.1109/TNSRE.2012.2200046.

